# Two forms of sexual dimorphism in gene expression in *Drosophila melanogaster*: their coincidence and evolutionary genetics

**DOI:** 10.1101/2021.02.08.429268

**Authors:** Amardeep Singh, Aneil F. Agrawal

**Affiliations:** Department of Ecology and Evolutionary Biology, University of Toronto, Toronto, Canada, M5S 3B2

## Abstract

Phenotypic sexual dimorphism can be mediated by sex differences in gene expression. We examine two forms of sexual dimorphism in gene expression in *Drosophila melanogaster*: (i) sex-biased gene expression (SBGE) in which the sexes differ in the amount a gene is expressed and (ii) sexual dimorphism in isoform usage, i.e., sex-specific splicing (SSS). In whole body (but not head) expression, we find a negative association between SBGE and SSS, possibly suggesting these are alternate routes to resolving sexual antagonistic selection. Next, we evaluate whether expression dimorphism contributes to the heterogeneity among genes in *r*_*mf*_, the intersexual genetic correlation in body expression that constrains the extent to which a gene’s expression can evolve independently between the sexes. We find lower *r*_*mf*_ values for genes with than without SSS. We find higher *r*_*mf*_ values for male- than female-biased genes (except genes with extreme male-bias), even though male-biased genes are known to have greater evolutionary divergence in expression. Finally, we examine population genetic patterns in relation to SBGE and SSS because genes with expression dimorphism have likely experienced a history of sex differences in selection. SSS is associated with reduced values of Tajima’s *D* and elevated *Direction of Selection* (*DoS*) values, suggestive of higher rates of adaptive evolution. Though *DoS* is highly elevated for genes with extreme male bias, *DoS* otherwise tends to decline from female-biased to unbiased to male-biased genes. Collectively, the results indicate that SBGE and SSS are differentially distributed across the genome and are associated with different forms of selection.

## Introduction

In sexual populations, the sexes employ different reproductive strategies to maximize fitness and as result, shared phenotypes often exhibit divergent, sex-specific trait optima. Because the sexes largely share a genome, the genetic basis of such traits is often the same between the sexes. Thus, genetic variation underpinning such shared traits will often experience divergent, sexually antagonistic selection resulting in an evolutionary conflict between the sexes termed ‘intra-locus sexual conflict’ (Arnqvist and Rowe 2005; Bonduriansky and Chenoweth 2009). Intra-locus sexual conflict has been well documented in a variety of plant (e.g., Hersh et al. 2015; Zemp et al. 2016; Delph et al.2022) and animal (e.g., Arnqvist and Rowe 2005; Cox and Calsbeek 2009) species and may be an important mechanism in maintaining genetic variation (Kidwell et al. 1977; Rice 1984). Ultimately, intra-locus sexual conflict is thought to be resolved through the evolution of sexual dimorphism (Lande 1980; Rice 1984).

Gene expression can be viewed as a trait in its own right or as a proximate mechanism affecting ‘traditional’ traits (e.g., morphological, behavioural, or physiological). Over the last ∼15 years, there has been a strong interest in investigating sexual dimorphism in gene expression, typically called ‘sex-biased gene expression’, SBGE (Ellegren and Parsch 2007; Parsch and Ellegren 2013; Mank 2017). SBGE refers to sex differences in the abundance of all transcripts mapping to a given gene. In addition to dimorphism in total expression of a gene, the sexes may differ in other aspects of the transcriptome. Here we examine SBGE as well as ‘sex-specific splicing’ (SSS), which here refers to quantitative differences in the relative usage of different isoforms between the sexes. Because the function of proteins and other gene products depends on the specific exons retained in mRNA transcripts after transcription, usage of different isoforms has the potential to greatly expand the functional repertoire of the genome, allowing a single gene to take on different functional roles across developmental stages (e.g., Gibilisco et al. 2016), tissue types (e.g., Barberan-Soler and Zahler 2008; Telonis-Scott et al. 2009; Revil et al. 2010; Ramani et al. 2011; Tan et al. 2013), and abiotic environments (Long et al. 2013; Jakšić and Schlötterer 2016).

Across the sexes, differential isoform usage (SSS) could allow the sexes to express their optimal trait values through the production of different gene products despite sharing a genome. Widespread SSS has been documented in humans (Trabzuni et al. 2013) and other primates (Blekhman et al. 2010) as well as in birds (Rogers et al. 2020). Several microarray and RNA-seq studies across *Drosophila* species have also shown that SSS is widespread (e.g., McIntyre et al. 2006; Telonis-Scott et al. 2009; Chang et al. 2011), tissue-specific (Telonis-Scott et al. 2009), and evolutionarily conserved (Gibilisco et al. 2016).

Here, we further explore SBGE and SSS using publicly available RNA-seq data from two tissue types (head and body tissue) of males and females of adult *Drosophila melanogaster*. As SBGE and SSS can represent different forms of sex-differential gene regulation, we are interested in their relationship. Resolution or mitigation of intra-locus sexual conflict could require the evolution of both SBGE and SSS; on the other hand, SSS and SBGE may serve as alternative routes to mitigating sexual conflict so that the evolution of one form of sex-differential regulation obviates selection for the other. We investigate whether genes that have one form of dimorphism are more or less likely to have the other, i.e., the co-occurrence of SSS and SBGE across the genome.

For any quantitative trait—including gene expression—a strong intersexual genetic correlation (*r*_*mf*_) constrains the potential for that trait to evolve independently between the sexes. Expression levels for most genes are positively genetically correlated across the sexes, yet the strength of the intersexual genetic correlation varies considerably among genes (Griffin et al. 2013; Allen et al. 2018). What factors contribute to this heterogeneity in *r*_*mf*_ values? It has been predicted that dimorphic traits will tend to have lower *r*_*mf*_ values than traits lacking dimorphism (Lande 1980; Bonduriansky and Rowe 2005). Using *Drosophila melanogaster*, Griffin et al. (2013) were the first to examine variation in gene expression *r*_*mf*_ values across the genome, finding—as predicted—that *r*_*mf*_ values tended to be lower in sex biased than unbiased genes. However, in their analysis, Griffin et al. (2013) did not distinguish on basis of the direction of the bias, i.e., male-versus female-biased genes. One might predict that male-biased genes would have lower *r*_*mf*_ values (Allen et al. 2018) given the observation that male-biased genes tend to diverge in their degree of sex bias more rapidly than female-biased genes (Meiklejohn et al. 2003; Zhang et al. 2007). With the benefit of a better dataset than was available to Griffin et al. (2013), we revisit these questions in *D. melanogaster*. Further, we ask if any of the heterogeneity in *r*_*mf*_ values among genes is related to SSS. Given that *r*_*mf*_ is expected to be reduced in traits with more dimorphism and that SSS represents another aspect of dimorphism in expression, we also expect *r*_*mf*_ would be lower, on average, in genes with SSS.

Genes with either form of expression dimorphism are likely to have experienced a history of differential selection between the sexes but the evolutionary consequences of differential selection are unclear. Compared to genes without dimorphic expression, are dimorphic genes more likely to experience balancing selection or relaxed selection or rapid adaptation? Because selection affects patterns of sequence diversity and divergence, comparing population genetic statistics among genes that differ in the strength, direction, and form of expression dimorphism may provide insights into selective forces acting on dimorphic genes (Meiklejohn et al. 2003; Pröschel et al. 2006; Fraïsse et al. 2019, Sayadi et al. 2019, Wright et al. 2019). For example, male-biased genes in *D. melanogaster* show signatures of higher rates of adaptive protein evolution than unbiased or female-biased genes (e.g., Pröschel et al. 2006; Fraïsse et al. 2018). We examine variation across the genome in three population genetic statistics—relative levels of nonsynonymous diversity, Tajima’s *D*, and direction of selection (*DoS*)—and test for associations of these with SBGE and SSS.

## Results and Discussion

### Prevalence of Sex-Specific Splicing

We find evidence of extensive sex-specific splicing (SSS) among multi-exon genes expressed in both head and body tissue of adult *Drosophila melanogaster*. Table 1 summarizes the results of our differential exon usage analysis. Of multi-exon genes tested, 5375 and 790 genes expressed in adult body and head tissue exhibit statistically significant SSS respectively which accounts for 57.5% and 10.2% of all genes tested in bodies and heads respectively. Considering the union of genes expressed in either tissue, we detected SSS in at least one of the tissues for 56% of genes. These data accord well with several previous studies that have found evidence of widespread SSS both in this species (e.g., McIntyre et al. 2006; Telonis-Scott et al. 2009; Chang et al. 2011) as well as other species of *Drosophila* (Gibilisco et al. 2016). McIntyre et al. (2006) reported that 68% of all multi-transcript genes expressed in whole body tissue exhibited significant sex-bias in the expression of one or more alternative isoforms in a microarray study using *D. melanogaster*. Similarly, a microarray study in this species by Telonis-Scott et al. (2009) found that ∼58% of multi-exon genes tested in whole body tissue exhibited SSS.

**Table 1:** Summary of the differential exon usage analysis between the sexes.

|  | Genes Tested | Genes with Sex-Specific Splicing<br>(% of genes tested) |
| --- | --- | --- |
| Head | 7772 | 790 (10.2%) |
| Body | 9346 | 5375 (57.5%) |
| <sup>1</sup>Head or Body | 9788 | 5488 (56.1%) |

<sup>1</sup>These data refer to the union of genes tested and exhibiting SSS found in either head, body or both.

Heads and bodies differ substantially in the frequency of SSS. Although the majority of genes exhibiting SSS in head tissue also exhibited SSS in the body, there are many genes exhibiting SSS in body tissue where we did not detect dimorphism in head tissue. 7330 genes were tested in both tissues. 4187 exhibit SSS in the body but only 677 (16%) of these also exhibit SSS in the head. Of the 7330, a total of 779 genes exhibit SSS in the head and 87% of these also exhibit SSS in the body. These results are consistent with the findings of Telonis-Scott et al. (2009) who reported reduced levels of SSS in genes when expressed in head tissue compared to carcass (i.e., head- and gonad-less bodies) and gonadal tissue.

### Coincidence of two forms of sexually dimorphism in expression

Numerous studies have examined sex differences variation in total expression for a gene. Studies of sex-biased gene expression (SBGE) have shown repeatedly that SBGE is highly tissue specific (e.g., Parisi et al. 2004; Yang et al. 2006; Dutoit et al. 2018); both our data and other studies (e.g., Telonis-Scott et al. 2009; Chang et al. 2011) indicate that the same is true for sex differences in isoform usage. Because both of these forms of sexual dimorphism are thought to create phenotypic dimorphism and help resolve intra-locus sexual conflict, we are interested in understanding the relationship between SBGE and SSS.

SBGE is widespread both in heads and bodies, though there is much greater variation among genes in the strength of SBGE in bodies relative to heads (e.g., Innocenti and Morrow 2010; Baker et al. 2011; Dutoit et al. 2018). To visualize the relationship between SSS and SBGE, we divided genes into bins representing different levels of sex-bias using defined ranges of log_2_FC in gene expression (see Methods). Within each bin, we calculated the proportion of genes exhibiting SSS. With respect to expression in the body, SSS is more common amongst unbiased genes than sex-biased genes, except for those with extreme sex-bias (Figure 1). This visual pattern is confirmed by the significantly negative quadratic SBGE term on the probability of SSS (*z* = -4.96; *p* = 7 × 10^−7^; Table S1). These fly body results are similar in this respect with Rogers et al. (2020) who reported non-significant trends towards a negative relationship between SSS and SBGE in gonadal tissue of three bird species.

**Figure 1.**
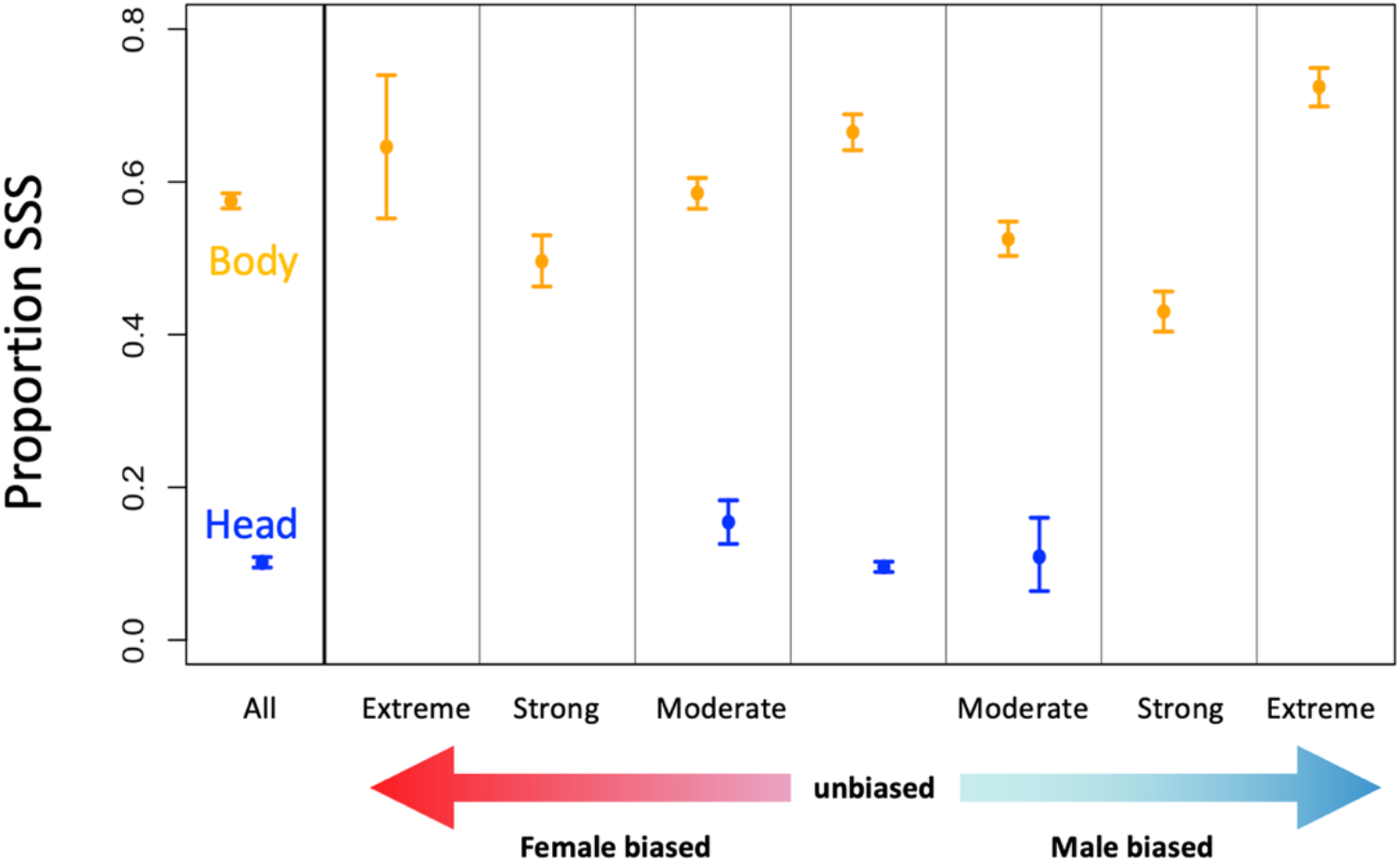
Proportion of genes exhibiting sex-specific splicing (SSS). The leftmost section shows results irrespective of SBGE; the remaining sections show results stratified by SBGE. Points in orange and blue represent reflect characterizations based on expression in body and head samples, respectively. Only three levels of SBGE are shown with respect to head because very few genes fall into the other levels. Error bars represent bootstrap 95% confidence intervals.

The reduction in isoform dimorphism at highly sex-biased genes expressed in the body could be because SBGE and SSS provide alternative routes to the resolution of sexual conflict and thus the evolution of one reduces the selective pressure for the evolution of the other. On the other hand, the pattern goes in the opposite direction with respect to expression in the head (Table S1, positive quadratic SBGE term, *z* = 3.02; *p* = 0.0025); SSS is less common among unbiased genes than sex-biased genes (at least those that are female-biased). One hypothesis for this latter pattern is that the resolution of intra-locus conflict may often require both forms of expression dimorphism whereas those genes experiencing sexually concordant selection will evolve neither, creating a positive association between SSS and SBGE. The difference in the relationships observed in head and body cannot be explained by differences in which genes are expressed. The patterns remain the same even if we consider only those genes expressed and tested for both SBGE and SSS in both tissues (Fig. S2).

An alternative, more mundane, explanation is that the positive association of SBGE and SSS in heads exists because of how we quantify gene/isoform expression (i.e., counting reads mapping to genes); we may be biased towards finding a positive association between SBGE and SSS because we are observing the same phenomenon in different ways. For example, if both males and females express a single isoform of a gene at the same level but females also express an additional isoform that is not expressed by males, we would detect both SBGE and SSS at this gene. However, this methodological bias must not be very strong as there are many unbiased genes that exhibit SSS (670 in head and 1007 in body). Moreover, the association between SBGE and SSS is negative in the body.

Given that the X chromosome spends a disproportionate amount of its time in females, and because of its hemizygous expression in males, there has been considerable interest in whether it is enriched for genes that experience sexually antagonistic selection contributing to intralocus conflict or contributing to sexual dimorphism (Rice 1987; Frank and Patten 2020). This has motivated interest in examining the location of genes exhibiting expression dimorphism. In the body data, there is no evidence of an enrichment of genes exhibiting SSS on the X chromosome (Figure 2A; permutation test: *p* < 0.78). In contrast, SBGE is strongly associated with the X chromosome, with female-biased genes being overrepresented on the X chromosome relative to unbiased genes whereas male biased genes are underrepresented on the X chromosome (Figure 2A; linear effect of SBGE on probability of being located on X: *z* = -7.06; *p* < 1.7 × 10^−12^). This observation of a “demasculinization” of the X chromosome has been well documented in previous studies of SBGE in *D. melanogaster* as well as several other species (Parisi et al. 2003; Ranz et al. 2003; Khil et al. 2004; Reinke et al. 2004; Sturgill et al. 2007; Meisel et al. 2012; Albritton et al. 2014), though Figure 2 illustrates the continuous, rather than discrete, nature of the relationship.

**Figure 2.**
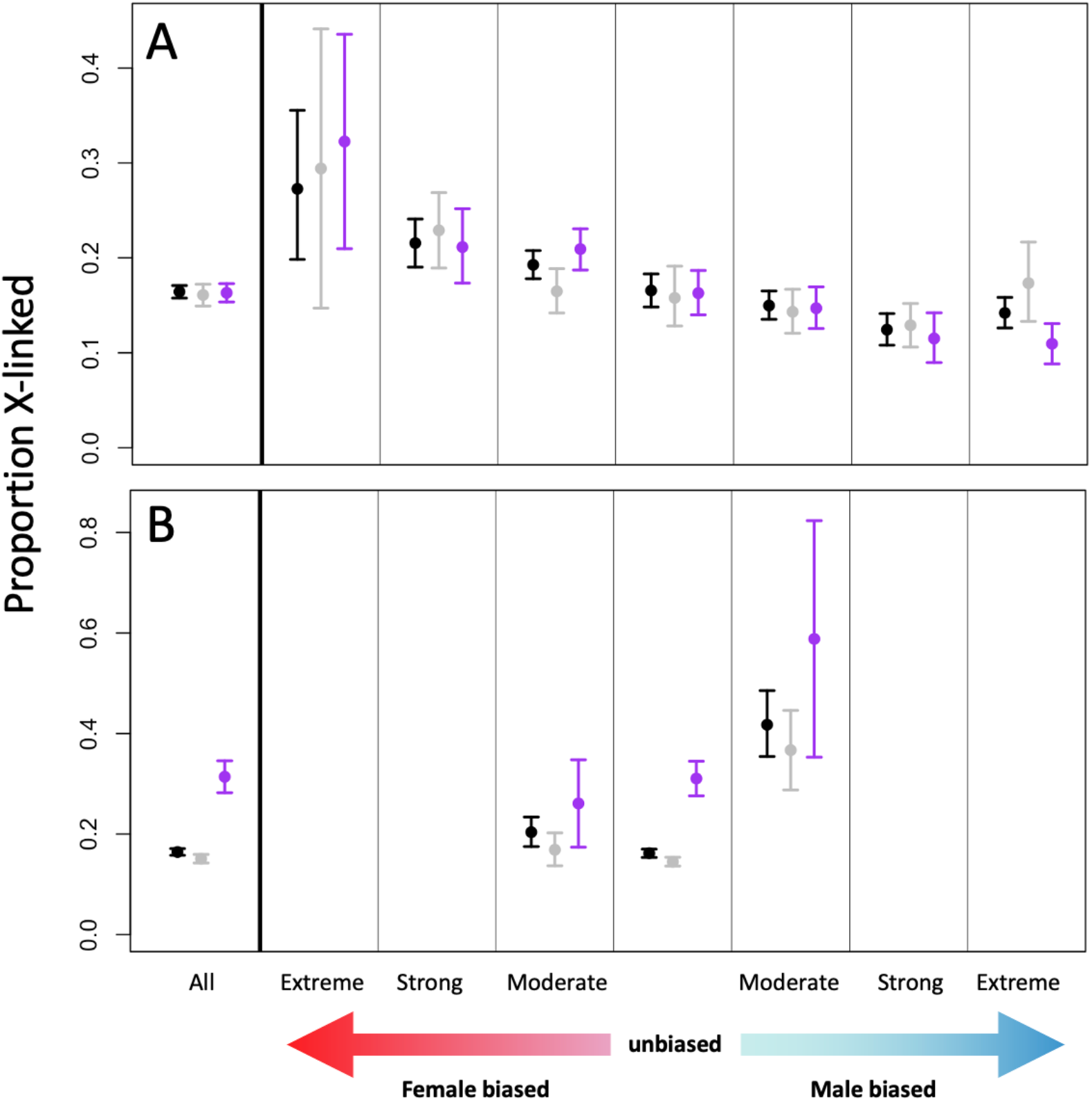
Proportion of X-linked genes with respect to SSS and SBGE as measured in (**A**) body and (**B**) head. The leftmost section in each panel shows results irrespective of SBGE; the remaining sections show results stratified by SBGE: extreme female bias (log_2_FC < -5), strong female bias (−5 ≤ log_2_FC < -2), moderate female bias (−2 ≤ log_2_FC < -0.5), unbiased expression (0.5 ≤ log_2_FC < 0.5), moderate male bias (0.5 ≤ log_2_FC < 2), strong male bias (2 ≤ log_2_FC < 5), and extreme male bias (log_2_FC ≥ 5). Points in grey and purple represent non-SSS and SSS genes, respectively; points in black are all genes irrespective of SSS status (including genes that were not tested for SSS so the black points represent more genes than the combined sum of genes represented by grey and purple points). Error bars are bootstrap 95% confidence intervals.

These patterns are quite different for the head data where there is a significantly higher proportion of X-linked genes exhibiting SSS relative to non-SSS genes (Figure 2B; permutation test of SSS vs non-SSS genes: *p* < 10^−4^). With respect to SBGE, both male- and female-biased genes are overrepresented on the X relative to unbiased genes, with the strongest overrepresentation among male-biased genes (Figure 2B; bootstrap 95% confidence intervals of proportion of X-linked genes for both male- and female-biased genes do not overlap with unbiased genes; see Meisel et al. (2012)). The over-representation of SSS genes on the X chromosome occurs primarily among unbiased genes rather than sex-biased genes, indicating that this pattern is not driven by a positive association between SSS and sex-biased expression.

### Association of SSS and SBGE with average expression

It is useful to consider how our two focal aspects of expression (SSS and SBGE) are associated with two other important aspects of expression—average expression intensity and tissue specificity—as each have been proposed or shown to correlate with population genetic indicators of selection (e.g., Osada 2007; Williamson et al. 2014; Fraïsse et al. 2018). We first consider a measure of the overall level of expression, measured as the log of FPKM averaged across a variety of tissues from larvae as well as adult males and females from the Fly Atlas 2 dataset (Krause et al. 2022). Average expression levels are higher for genes with versus without SSS (Fig. 3; permutation test: *p* < 10^−4^ for both body and head). To some extent, this will be due to the greater power to detect SSS in genes that have higher expression. Regardless of the underlying reason, this association will be an important consideration when we later consider evolutionary characteristics of genes with SSS. In bodies, average expression is greater for unbiased than sex-biased genes (quadratic effect of SBGE on average expression is significantly negative: *z* = -33.6, *p* < 2 × 10^−16^). In heads, average expression tends to decline as we move from female-biased genes to unbiased genes to male-biased genes (linear effect of SBGE on average expression is significantly negative: *z* = -22.9, *p* < 2 × 10^−16^).

**Figure 3.**
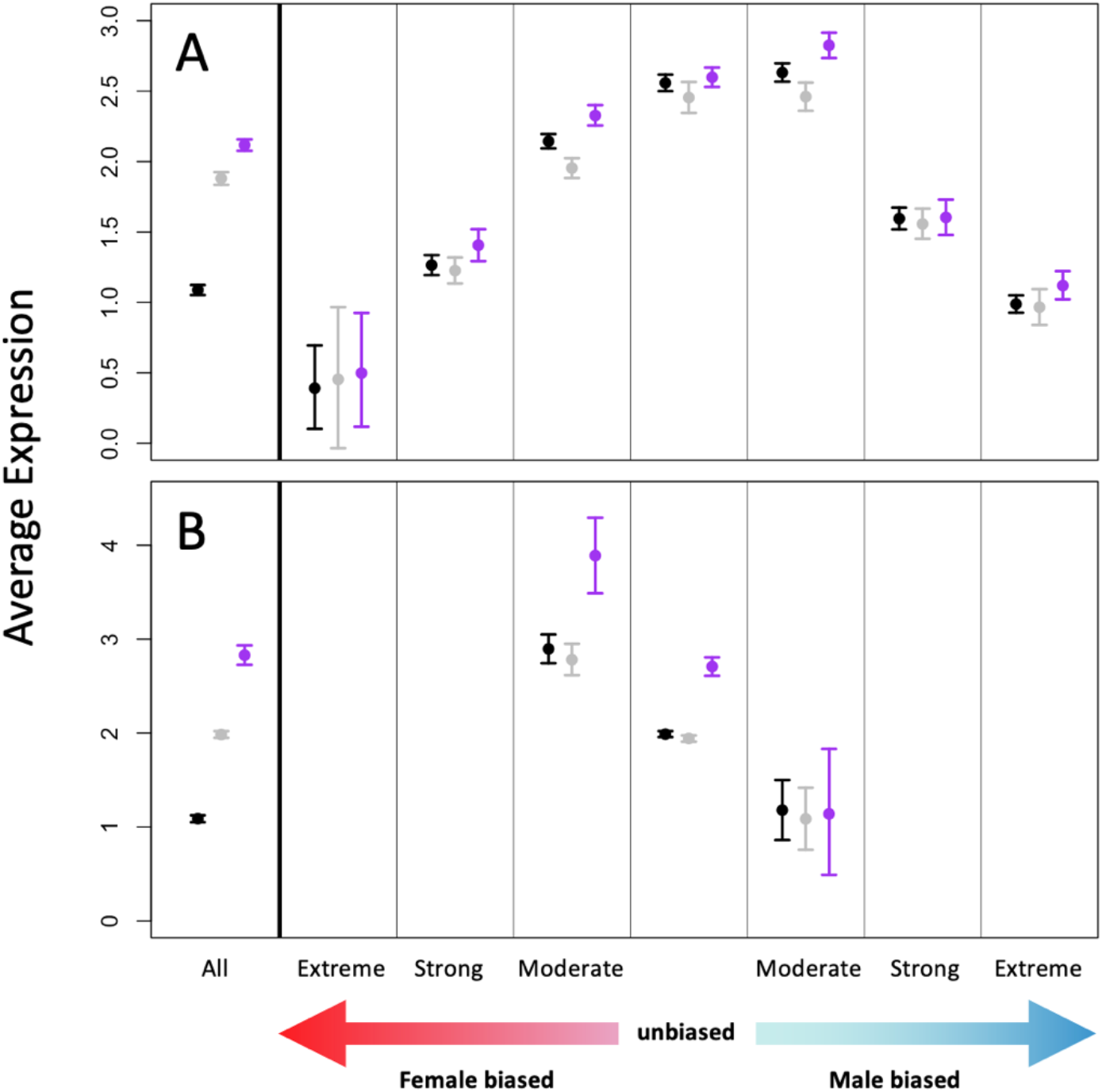
Association of SSS and SBGE with average expression. Expression level averaged across multiple tissues in both larval and adult tissues is shown with respect to SSS and SBGE in (**A**) body and (**B**) head. The leftmost section in each panel shows results irrespective of SBGE; the remaining sections show results stratified by SBGE: extreme female bias (log_2_FC < -5), strong female bias (−5 ≤ log_2_FC < -2), moderate female bias (−2 ≤ log_2_FC < -0.5), unbiased expression (0.5 ≤ log_2_FC < 0.5), moderate male bias (0.5 ≤ log_2_FC < 2), strong male bias (2 ≤ log_2_FC < 5), and extreme male bias (log_2_FC ≥ 5). Points in grey and purple represent non-SSS and SSS genes, respectively; points in black are all genes irrespective of SSS status (including genes that were not tested for SSS so the black points represent more genes than the combined sum of genes represented by grey and purple points). Error bars are bootstrap 95% confidence intervals.

### Association of SSS and SBGE with tissue specificity

The other aspect of expression we consider is tissue specificity, based on the evenness of expression across multiple tissues in larva as well as adult males and females calculated using data from Fly Atlas 2 (Krause et al. 2022). Tissue specificity is higher for sex biased than unbiased genes (Fig. 4; quadratic effect of SBGE on tissue specificity is significantly positive: body *z* = 51.2, *p* < 2 × 10^−16^; head *z* = 26.5, *p* < 2 × 10^−16^). Tissue specificity is lower for SSS than non-SSS genes (permutation test SSS vs non-SSS genes: body *p* < 10^−4^; head *p* < 10^−4^).

**Figure 4.**
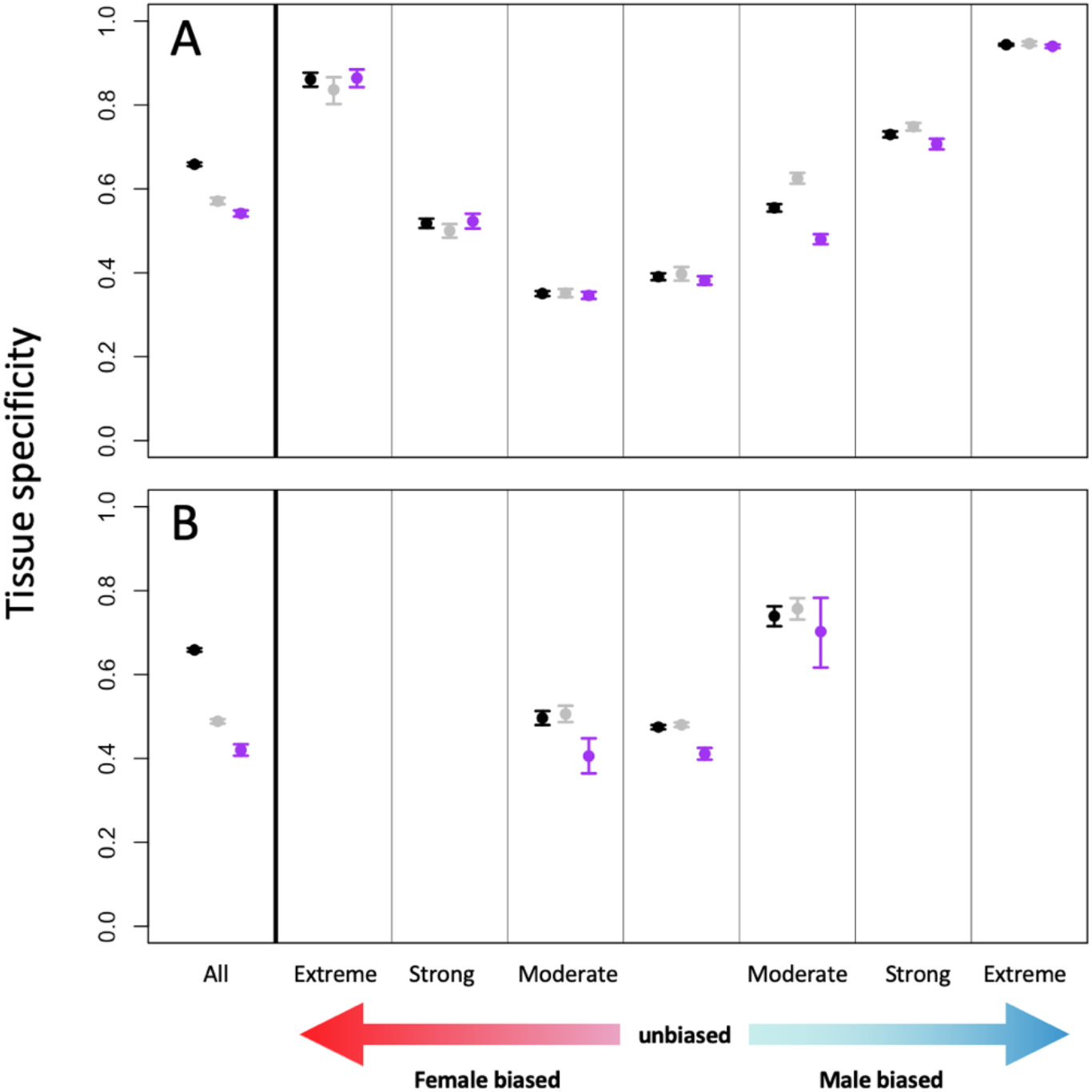
Mean tissue-specificity with respect to SSS and SBGE as measured in (**A**) body and (**B**) head. The leftmost section in each panel shows results irrespective of SBGE; the remaining sections show results stratified by SBGE: extreme female bias (log_2_FC < -5), strong female bias (−5 ≤ log_2_FC < -2), moderate female bias (−2 ≤ log_2_FC < -0.5), unbiased expression (0.5 ≤ log_2_FC < 0.5), moderate male bias (0.5 ≤ log_2_FC < 2), strong male bias (2 ≤ log_2_FC < 5), and extreme male bias (log_2_FC ≥ 5). Points in grey and purple represent non-SSS and SSS genes, respectively; points in black are all genes irrespective of SSS status (including genes that were not tested for SSS so the black points represent more genes than the combined sum of genes represented by grey and purple points). Error bars are bootstrap 95% confidence intervals.

Previous studies in *Drosophila* (Assis et al. 2012; Fraïsse et al. 2019) and other animals (Mank et al. 2008; Dean and Mank 2016) have also found that sex-biased genes have higher tissue specificity than unbiased genes; this has been interpreted as sexual dimorphism evolving more readily when there is less pleiotropic constraint on expression. Under this premise, one might expect the same for SSS. However, in the body data, SBGE is associated with greater tissue specificity whereas SSS is associated with lower tissue specificity. Similarly, Rogers et al. (2020) found that genes exhibiting SSS exhibited lower tissue specificity in gene expression than non-SSS genes in birds.

### Associations of SSS and SBGE with Evolutionary Properties

We next examine whether SSS and SBGE are related to one quantitative genetic property (*r*_*mf*_) and three population genetic properties (relative nonsynonymous diversity, Tajima’s *D*, and *DoS*). We do this using SSS and SBGE as measured in the body, not head. The available data on *r*_*mf*_ is based on whole body expression so it is appropriate to analyze variation in *r*_*mf*_ in relation to body-based measures of SSS and SBGE. The analysis of the population genetic properties is motivated by the idea that genes with dimorphic expression are more likely than non-dimorphic genes to have a history of sex differences in selection; analysis of the population genetic properties may reveal insights into the nature of this selection. Our *a priori* expectation is that dimorphism as measured from body, rather than head, samples is a better indicator of the likelihood that a gene has experienced a history of differential selection between the sexes. For example, many genes have dimorphic expression in body but not head and it is likely more reasonable for these analyses to characterize those genes as dimorphic (and likely to have experienced a history of differential selection between the sexes) than as non-dimorphic.

For each evolutionary genetic property, we examine associations with SSS and SBGE in two ways. First, we examine simple associations of each evolutionary property with SSS and SBGE, which are straightforward to represent graphically. This approach ignores other factors that could be important sources of variation in the focal evolutionary property. Such factors could confound or obscure a relationship with SSS or SBGE if they covary with SSS or SBGE. Consequently, we also use linear models in which we examine variation in an evolutionary property as a function of SSS and SBGE as well as other relevant covariates. These other covariates include X-linkage (i.e., X or autosomal) as well as several other aspects of expression: average expression, tissue specificity, and summary measures of the across-tissue expression profile for each sex. The latter are the values from the three leading principal components (PCs) of variation in across-tissue expression profiles within each sex (i.e., 6 PCs total). In the analyzes of the population genetic properties, we also included two other covariates that could affect the strength of linked selection: recombination rate and gene length. As we show graphically, genes with extreme sex bias (particularly extreme male-bias) often differ dramatically from other genes. We exclude these genes from all the linear models. The vast majority of genes that could be regarded as “sex-limited”, i.e., having very low read counts in one sex, fall into the extreme sex-bias categories (especially the extreme male-bias category). Such genes may only experience selection in only one sex and this may be an important reason for their unique properties.

### *Intersexual genetic correlation*, r_mf_

The extent to which a gene’s expression level can evolve independently in one sex from the other is mediated by the intersexual genetic correlation in expression level, *r*_*mf*_. Genes with dimorphic expression are predicted to have reduced *r*_*mf*_. (This could be either because dimorphism evolves more easily for genes with low *r*_*mf*_ or because the process of evolving dimorphic expression results in an increased degree of independence of how expression is regulated between the sexes, i.e., lower *r*_*mf*_ evolves as a by-product of increased expression dimorphism.) Previous work in *D. melanogaster* found that *r*_*mf*_ values decline moving from unbiased to increasingly sex-biased genes but did not distinguish between male- and female-biased genes (Griffin et al. 2013). This distinction is important as male-biased genes diverge in their extent of SBGE more rapidly female-biased genes (Meiklejohn et al. 2003; Zhang et al. 2007) and one potential contributing factor could be a lower *r*_*mf*_ in male-biased genes (Allen et al. 2018).

We find a striking relationship between *r*_*mf*_ and SBGE, with female-biased genes having lower *r*_*mf*_ than unbiased genes (non-overlapping 95% confidence intervals in Fig. 5A). Moderately and strongly male-biased genes have the highest average *r*_*mf*_ values. In contrast, genes with extreme male bias have very low *r*_*mf*_ values on average. A linear model that helps control for correlated variables indicates that *r*_*mf*_ increases going from female-biased to unbiased to male-biased genes (significant linear effect of SBGE in Table 2; note this analysis excludes genes with extreme sex bias). In *Drosophila serrata, r*_*mf*_ is also higher in male-than female-biased genes (Allen et al. 2018). These results imply that the faster divergence in SBGE in male-than female-biased genes occurs despite the higher *r*_*mf*_ values of the former, presumably due to a higher rate in changes in selection on expression of male-than female-biased genes.

**Figure 5.**
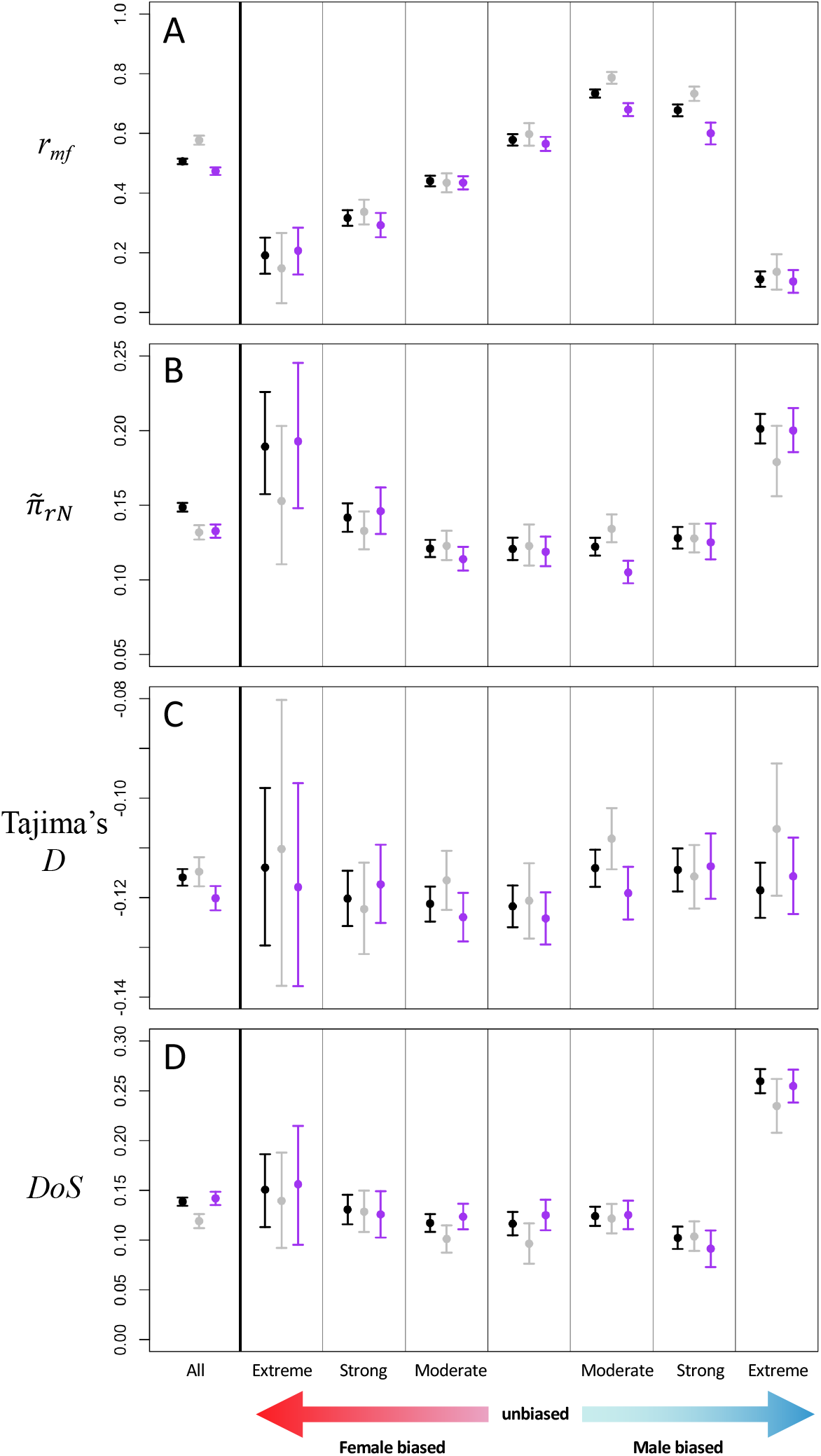
Four evolutionary genetic properties in relation to expression properties in the body. The leftmost section in each panel shows results irrespective of SBGE; the remaining sections show results stratified by SBGE: extreme female bias (log_2_FC < -5), strong female bias (−5 ≤ log_2_FC < -2), moderate female bias (−2 ≤ log_2_FC < -0.5), unbiased expression (0.5 ≤ log_2_FC < 0.5), moderate male bias (0.5 ≤ log_2_FC < 2), strong male bias (2 ≤ log_2_FC < 5), and extreme male bias (log_2_FC ≥ 5). Points in grey and purple represent non-SSS and SSS genes, respectively; points in black are all genes irrespective of SSS status (including genes that were not tested for SSS so the black points represent more genes than the combined sum of genes represented by grey and purple points). Error bars are bootstrap 95% confidence intervals.

**Table 2.** Results from linear models examining genome-wide variation in *r*_*mf*_, 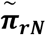, Tajima’s *D* and *DoS*.

| Model Term | $r_{mf}$ | $\tilde{\pi}_{rN}$ | Tajima's $D$ | $DoS$ |
| --- | --- | --- | --- | --- |
| SSS | **-0.014\*\***<br>[-0.024, -0.004] | 0.001<br>[-0.003, 0.005] | **-0.003\***<br>[-0.006, -0.0001] | **0.011\*\*\***<br>[0.005, 0.016] |
| SBGE | **0.041\*\*\***<br>[0.034, 0.048] | -0.002<br>[-0.005, 0.001] | 0.000<br>[-0.002, 0.003] | **-0.006\*\***<br>[-0.011, -0.002] |
| SBGE<sup>2</sup> | **-0.006\*\*\***<br>[-0.009, -0.003] | 0.000<br>[-0.001, 0.001] | 0.000<br>[-0.0001, 0.0004] | 0.000<br>[-0.001, 0.001] |
| Avg Expression | **0.057\*\*\***<br>[0.050, 0.065] | **-0.011\*\*\***<br>[-0.014, -0.007] | **-0.004\*\*\***<br>[-0.007, -0.002] | **0.011\*\*\***<br>[0.005, 0.016] |
| Tissue Specificity | **0.250\*\*\***<br>[0.198, 0.300] | **0.051\*\*\***<br>[0.029, 0.073] | **0.027\*\*\***<br>[0.011, 0.042] | 0.008<br>[-0.026, 0.040] |
| M-PC1 | **0.044\*\***<br>[0.037, 0.051] | -0.002<br>[-0.005, 0.0005] | **0.003\*\*\***<br>[0.001, 0.005] | **-0.012\*\*\***<br>[-0.016, -0.008] |
| M-PC2 | **0.018\*\*\***<br>[0.009, 0.026] | 0.001<br>[-0.002, 0.004] | **0.003\*\***<br>[0.001, 0.006] | -0.001<br>[-0.006, 0.003] |
| M-PC3 | **0.021\*\*\***<br>[0.014, 0.027] | 0.001<br>[-0.002, 0.003] | 0.000<br>[-0.002, 0.002] | -0.003<br>[-0.007, 0.001] |
| F-PC1 | 0.003<br>[-0.006, 0.011] | 0.001<br>[-0.002, 0.004] | 0.000<br>[-0.002, 0.002] | -0.002<br>[-0.007, 0.001] |
| F-PC2 | **0.013\*\*\***<br>[0.006, 0.019] | **0.003\***<br>[0.001, 0.006] | -0.001<br>[-0.003, 0.001] | 0.001<br>[-0.003, 0.005] |
| F-PC3 | -0.002<br>[-0.007, 0.003] | 0.001<br>[-0.002, 0.003] | 0.000<br>[-0.002, 0.001] | 0.001<br>[-0.002, 0.004] |
| X-linkage | **0.026\*\*\***<br>[0.015, 0.038] | **-0.007\*\***<br>[-0.012, -0.002] | **0.021\*\*\***<br>[0.018, 0.025] | **-0.030\*\*\***<br>[-0.037, -0.022] |
| log(recombination) | - | **-0.021\*\*\***<br>[-0.023, -0.018] | **-0.002\*\***<br>[-0.004, -0.001] | **0.022\*\*\***<br>[0.019, 0.025] |
| log(gene length) | - | **-0.022\*\*\***<br>[-0.028, -0.016] | **0.006\*\***<br>[0.002, 0.010] | **-0.027\*\*\***<br>[-0.036, -0.018] |

The numbers of genes used for the four analyses are $n = 4982, 4624, 4750$ , and $4761$ . For these analyses, genes with extreme sex-bias ( $\log_2FC > |5|$ ) are excluded. The quadratic term, SBGE<sup>2</sup>, is included to capture effects of sex bias irrespective of the direction of the bias. Estimates in bold have significant permutation-based $p$ -values: \* $p < 0.05$ , \*\* $p < 0.01$ , \*\*\* $p < 0.001$ . 95% bootstrap confidence intervals are shown in square brackets.

For SSS to exist, there must already be some sex differences in how gene expression is regulated. Thus, one might further predict that expression levels across sexes for genes with SSS are less likely to be strongly genetically correlated. Consistent with this prediction, we find that *r*_*mf*_ is lower, on average, for genes with vs. without SSS (permutation test: *p* < 10^4^; Fig. 5A). This effect of SSS remains significant in the linear model with other covariates, including SBGE (Table 2). These results indicate that expression levels in the two sexes should be able to evolve more independently in genes with SSS. This generates a prediction for future studies that genes with SSS should tend to show higher rates of divergence in SBGE, assuming all else equal.

Other aspects of expression are also significantly associated with *r*_*mf*_ (Table 2). *r*_*mf*_ is higher for genes with higher average expression and genes with higher tissue specificity. The latter result seems counter intuitive because one might expect a greater degree of genetic independence between the sexes (i.e., lower *r*_*mf*_) for genes expressed predominantly in highly dimorphic or sex-limited tissues (e.g., gonads, accessory glands, spermatheca). However, it is worth remembering the analysis in Table 2 excludes genes with extreme sex bias and thereby excludes many genes with high specificity to such tissues. Among the remaining genes, genes with high tissue specificity will tend to be predominantly expressed in tissues that are typically less dimorphic (e.g., heart, midgut, tubule, etc.) whereas the genes with low tissue specificity will include genes that are expressed both in highly dimorphic tissues (e.g., gonads) as well as in other tissues. This may explain the surprising positive association of tissue specificity with *r*_*mf*_. In addition, *r*_*mf*_ is associated with aspects of the across-tissue expression profile in both sexes (as captured by the PC terms). This indicates that genes with similar across-tissue expression profiles tend to have similar *r*_*mf*_ values, possibly reflecting the effects of allometry and tissue heterogeneity on whole body expression measures. The direction of associations of *r*_*mf*_ with the specific PCs appear consistent with our preceding comments. For example, *r*_*mf*_ is positively associated with male PC1 and with female PC2; the loading for testis expression is strongly negative on M-PC1 and the loading for ovary expression is strongly negative on F-PC2. These patterns indicate that *r*_*mf*_ is lower in genes with greater gonad expression, matching intuition. Though these associations of *r*_*mf*_ with other aspects of expression are interesting, our focus here is on SSS and SBGE. Within the limits of inference from linear modeling, our analysis indicates that SSS and SBGE are significantly associated with *r*_*mf*_ even after accounting for these other aspects of expression.

### Relative level of nonsynonymous diversity, 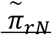

A reduction in nucleotide diversity for nonsynonymous relative to synonymous variants is typically interpreted as indicative of purifying selection on amino acid sequence variation, with lower nonsynonymous diversity levels indicating stronger selection. We calculated 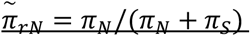 from 199 genotypes from the DPGP3 (Lack et al. 2015, 2016). A simple inspection of relative levels of nonsynonymous diversity 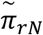 reveals no clear difference between genes with versus without SSS (Fig. 5B). Genes with extreme sex-bias have elevated levels of 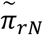_;_ (i.e., non-overlapping 95% confidence intervals in Fig. 5B), consistent with the idea that such genes experience weaker net purifying selection because they are primarily selected in only one sex (Pröschel et al. 2006; Dapper and Wade 2016, 2020). In the linear model, for which we exclude the extreme sex-bias genes, there are no significant effects of SBGE or SSS.

### *Tajima’s* D

Tajima’s *D* is a summary statistic that reflects information about the shape of the underlying genealogy, which can be affected by the strength and form of selection at linked sites. As such, variation in Tajima’s *D* has been used to look for indirect evidence that genes with different expression properties are selected differently (Sayadi et al. 2019; Wright et al. 2019). As shown in Fig. 5C, genes with SSS have lower values of Tajima’s *D* (permutation test: *p* < 0.007). There is no visually obvious relationship of Tajima’s *D* with SBGE (Fig. 5C). The results of the linear model are consistent with these findings for both SSS and SBGE (Table 2).

The result that SSS and non-SSS genes differ with respect to Tajima’s *D* suggests these two sets of genes differ with respect to some aspect of selection. There is no evidence that there is a consistent difference in the strength of purifying selection given that SSS and non-SSS genes do not differ with respect to 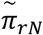. This leaves open the possibility that the difference in Tajima’s *D* could be because genes with SSS are less likely to experience balancing selection or more likely to experience selective sweeps than genes without SSS.

### *Direction of Selection* DoS

An excess of divergence to polymorphism for nonsynonymous relative to synonymous variants can be indicative of adaptive evolution (McDonald and Kreitman 1991). Based on this idea, Stoletzki and Eyre-Walker (2011) devised the *DoS* metric for which positive values are indicative of adaptive evolution, with more strongly positive values reflecting more frequent adaptive evolution. *DoS* is higher for genes with SSS than those without (permutation of SSS vs non-SSS gene: *p* < 0.006; see also Table 2). With respect to SBGE, *DoS* is substantially elevated for genes with extreme male-bias (Fig. 5D). Excluding the genes with extreme sex-bias, the linear model detects a subtle but significant effect of SBGE on *DoS* (Table 2), indicating a decline in *DoS* going from female-biased to unbiased to male-biased genes.

A potential concern in interpretating *DoS* is that large positive values could be primarily due to low levels of nonsynonymous polymorphism, with divergence playing little role. However, divergence and polymorphism components each contribute substantially to variation in *DoS* in this dataset. Moreover, if we apply a linear model to just the divergence component of *DoS* (i.e, *D*_*N*_*/*(*D*_*N*_ + *D*_*S*_)), we again detect a positive effect of SSS (Table S2). As with *DoS*, the divergence component of *DoS* declines from female-biased to unbiased to male-biased genes (Table S2; this analysis again excludes genes with extreme sex bias, genes with extreme male-bias have highly elevated values for the divergence component of *DoS*, Fig. S4).

### Other aspects of expression

Though our focus is on SSS and SBGE, we note that each of the population genetic metrics have significant associations with other aspects of expression (Table 2). For example, the linear model indicates 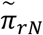 is lower for more highly expressed genes. This is consistent with results from a variety of taxa, which is usually interpreted as indicating stronger purifying selection on highly expressed genes (e.g., Carneiro et al. 2012; Paape et al. 2012; Williamson et al. 2014; Ho et al. 2019). Though average expression is also associated with higher values of *DoS*, this is likely a reflection of purifying selection rather than adaptation as average expression has a significantly negative association with the polymorphism component of *DoS* but is not associated with the divergence component of *DoS* (Table S2, Fig. S4; see also Fraïsse et al. 2019 and Huang 2022). The analysis of 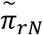 also indicates that higher tissue specificity results in higher values of 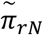, consistent with the idea of weaker purifying selection on genes with less pleiotropy (e.g., Osada 2007; Paape et al. 2013).

## Conclusions and further considerations

SSS and SBGE are two different ways to differentially regulate genes across the sexes and thus could be mechanisms by which sexual conflict is resolved. Genes with SSS have lower values of the intersexual genetic correlation in expression *r*_*mf*_. This indicates that patterns of standing genetic variation are more conducive to evolving greater sex differences in a gene’s total expression (i.e., stronger SBGE) for genes with SSS versus without, holding the amount of genetic variation constant. Thus, one might predict genes with SSS to have higher levels of SBGE, which stands at odds with the observed negative association between these two forms of expression dimorphism in the body (Fig. 1). This could be because when one of these mechanisms evolves, it may negate the selective pressure to evolve the other (i.e., SSS genes are less likely to be under antagonistic selection for total expression than non-SSS genes).

SSS and SBGE are each associated with some of the population genetic metrics we examined, suggesting that genes with expression dimorphism tend to experience somewhat different selection than non-dimorphic genes. Further, there are some differences between SSS and SBGE in how each are associated with population genetic metrics, suggesting differences in selection between genes with these two different forms of dimorphism. For example, SSS, but not SBGE, is associated with a reduction in Tajima’s *D*. An interpretation of the analyses Tajima’s *D* and *DoS* together is that SSS genes experience more adaptive evolution than genes without SSS. Perhaps when sexes use different isoforms, it is easier for adaptation to occur in each sex because there is less pleiotropic constraint imposed by the other sex. In contrast, the pattern with respect to SBGE (excluding genes with extreme sex-bias) is less clear. The *DoS* analysis does not imply sex-biased genes experience more adaptive evolution than unbiased genes, though this may be because adaptive evolution is modestly elevated among female-biased genes but modestly reduced among male-biased genes relative to unbiased genes.

Such interpretations should be regarded with caution for several reasons. First, population genetic metrics can be influenced by a multitude of factors and the interpretations above are not necessarily correct. Second, the comparison of SSS and SBGE is complicated by the nature of how they are defined and measured. SBGE has both a direction (i.e., negative to positive representing female-to male-biased) and a magnitude. SSS, as measured here, simply indicates whether the sexes differ in their distribution of isoform usage; there is no male/female direction. Magnitude only affects SSS status via the statistical power to detect the difference, which also depends on other factors (e.g., expression level, sequencing depth). (Quantifying a “magnitude” for SSS is complicated by the multitude of ways that isoform distributions can differ; but see Rogers et al. 2020.)

An additional reason for caution is an unmeasured correlated variable could be responsible for the reported associations of evolutionary properties with SBGE and SSS. For this reason, we incorporated several plausibly important variables into our analyses to reduce this possibility. One aspect of this warrants further discussion. Whole body samples represent a collection of tissue types and expression profiles differ across tissues. Thus, expression differences between sexes can be due to either expression changes within tissues (or cell types) between sexes or the relative contribution of each tissue within each sex (Stewart et al. 2010). We take the perspective that expression in whole body samples represents a coarse-grained level at which dimorphism can be studied, while recognizing that several different biological levels could contribute to this level of dimorphism. Examining dimorphism at this level we find that SBGE and SSS are associated with several key evolutionary properties. A potential concern is that these associations are due to some other feature of genes that is associated with these measures of expression dimorphism. For example, perhaps genes predominantly expressed in particular tissues (e.g., gonads) evolve differently than other genes expressed predominantly in other tissues (e.g., midgut) but genes in the former tissues tend to have different levels of expression dimorphism measured at the whole body level than other genes (i.e., perhaps SBGE and SSS are just crude indicators of a gene’s among-tissue expression profile either because particular tissues have more strongly dimorphic expression or because the relative sizes of particular tissues differs considerably between sexes).

To mitigate this concern, our linear models explicitly included principal components (PCs) reflecting variation in across-tissue expression profiles in both sexes. For every population genetic metric we examined, one or more of the PC terms was significant, suggesting that genes with similar across-tissue expression profiles experience, to some extent, similar population genetic pressures. However, the key point with respect to our primary interest is that the effects of SBGE and SSS are significant even after accounting for variation in across-tissue expression profiles. Further, statistical models without the PC terms yield similar estimates for the effects of SBGE and SSS, providing no indication that considering variation in across-tissue expression profiles has a substantial influence on the inferred effects of SBGE and SSS. While it would be naïve to presume that the inclusion of the PCs in the analysis controls for all possible tissue effects, our analyses show SBGE and SSS are associated with population metrics in ways that not readily explained by tissue effects. Thus, the observed patterns with respect to SBGE and SSS are intriguing; an on-going challenge is to better understand why they occur.

## Methods

### Obtaining RNA-seq Data and Read Mapping

We obtained publicly available raw RNA-seq data from the DNA Data Bank of Japan Sequence Read Archive (Accession number: DRA002265). Detailed methods on animal husbandry, RNA extraction and sequencing can be found in Osada et al. (2017). Briefly, males from 18 randomly chosen inbred lines of the *Drosophila* Genomic Resource Panel (DGRP; MacKay et al. 2012; Huang et al. 2014) were crossed to females from a single inbred strain from a West African population. RNA was extracted from heads and bodies (i.e., whole body minus heads) of 100 virgin F1 males and females from each cross. RNA from female heads and bodies was obtained and sequenced from two replicates of each of the 18 DGRP lines while only a single replicate was available for males. We therefore arbitrarily retained sequence data only for female samples labelled as ‘replicate 1’. Reads were inspected for quality using *FastQC* (v0.11.9; Andrews 2010) and *MultiQC* (v0.8.3; Ewels et al. 2016). Reads were trimmed for adapter content using *Trimmomatic* (v0.39; Bolger et al. 2014). RNA-seq reads were then mapped to the *Drosophila melanogaster* reference genome (release 6.32; Dos Santos et al. 2015) using *STAR* (v2.7.1a; Dobin et al. 2013) with default settings.

### Sex-Specific Splicing

To identify a list of genes exhibiting SSS, we performed differential exon usage analysis using the R package *JunctionSeq* (v1.17; Hartley and Mullikin 2016). Briefly, *JunctionSeq* splits genes into annotated exons and splice junction ‘counting bins’ and counts the number of reads that overlap each counting bin within an annotated gene. To test for differential exon usage between the sexes, a generalized linear model (GLM) with logarithmic link function was fit at each gene feature (i.e., exon or splice junction) to assess sex differences in the number of reads that map to a particular gene feature relative to the total number of reads that map to the gene to which the feature belongs. Specifically, for each gene, *JunctionSeq* fits the model

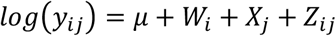

where *y*_*ij*_ is the normalized count of reads from the *i*^th^ sex mapping to the *j*^th^ gene feature/counting bin. *µ* is the experiment wide, mean normalized gene expression, *W*_*i*_ reflects the expected number of reads mapping to the focal gene in sex *i, X*_*j*_ reflects the proportion of reads mapping to counting bin *j*, and*Z*_*ij*_ is an interaction term that reflects differences between the sexes in the proportion of reads mapping to counting bin *j*. A significant effect of the interaction term was taken as evidence of SSS. To test the significance of the interaction term, the above model was compared to a null model that was identical, but which omits the interaction term.

*JunctionSeq* can test genes with at least two annotated exons. It applies a multiple-testing correction method based on the Benjamini and Hochberg procedure (Benjamini and Hochberg 1995) to generate adjusted *P-*values. Genes were considered to show evidence for sexual dimorphism in isoform usage if the sexes exhibited statistically significant differential usage of a single exon (or more) within a gene using *p*_*adj*_ < 0.05.

Ideally, genotype would be included in the model as a random effect but *JunctionSeq* does not allow this. This is unlikely to be a major issue here. Our dataset is perfectly balanced with each genotype being represented by only a single sample for each sex for each tissue (head and body data were analyzed separately). More importantly, our goal is to identify a list of genes enriched for SSS for further analysis; our focus is not on the accuracy of *p*-values of individual genes. As a simple check of the *JunctionSeq* pipeline, we analyzed 10 datasets in which we switched the male/female labels for a randomly chosen half of the genotypes. In these permuted data sets, no genes were identified as having SSS at *p*_*adj*_ < 0.01 whereas thousands of genes are found in the true (unpermuted) data under the same criterion.

### Sex-biased gene expression

We estimated differential gene expression between the sexes using the R package *DESeq2* (v1.28.1; Love et al. 2014). Briefly, *DESeq2* estimates normalized read counts mapping to genes on a log_2_ scale and uses a GLM framework to test for differential gene expression between experimental conditions. Before performing differential gene expression analysis, we performed a ‘pre-filtering’ procedure to remove genes that had very low numbers of reads mapping. Pre-filtering RNA-seq data has been shown to substantially increase power in detecting differential gene expression by reducing the burden of multiple testing but can also result in an increase in Type I error (i.e., increased false-positive rate) (Bourgon et al. 2010). To balance these competing issues, we removed any genes that had fewer than an average of 50 reads mapping from all 36 samples (i.e., 18 male and 18 female genotypes). We then estimated SBGE as the log_2_ fold change (log_2_FC) in male to female gene expression for each annotated gene and retained only genes that were tested for differential exon usage. In downstream analyses, we assigned each gene to bins based on based on the log2FC in male to female gene expression.

For visual inspection of patterns, we assigned each gene to one of seven bins of predefined ranges of log_2_FC in gene expression: extreme female bias (log_2_FC < -5), strong female bias (−5 ≤ log_2_FC < -2), moderate female bias (−2 ≤ log_2_FC < -0.5), unbiased expression (0.5 ≤ log_2_FC < 0.5), moderate male bias (0.5 ≤ log_2_FC < 2), strong male bias (2 ≤ log_2_FC < 5), and extreme male bias (log_2_FC ≥ 5). Binning was conducted separately with respect to expression in heads and bodies. A small number of genes for which the standard error in log_2_FC was high (> 0.5) were excluded. The bins were used for graphical presentation of results; as described below, our main analyses treat log_2_FC as a continuous variable. In heads, there were few genes with log_2_FC > |2| so only the three central bins are shown in the figures. Distributions of log_2_FC for each tissue are shown in Fig. S1.

Some genes could be regarded as “sex-limited” based on very low expression in one sex, though there is no universally accepted definition. Operationally defining “sex-limited” as an average read count across samples below 5 in only one sex, almost all such genes fall into the extreme sex-bias categories (body: 959/963 = 99.6% [17 and 942 genes are extreme female- and male-biased, respectively], head: 2/2 = 100% [both are extreme female-biased]). Not all genes classified as having extreme sex-bias in the body would be classified as sex-limited under this (arbitrary) definition; 79/96 = 82% of extreme female-biased genes and 228/1170 = 19% of extreme male-biased genes would not qualify as “sex-limited” under this definition even though the vast majority of genes with extreme sex bias have low to very low expression in one sex. No analyses are based upon the classification of “sex-limited”. Rather, we are simply alterting readers that the “extreme sex-bias” categories include many genes that may be regarded as having sex-limited expression.

### Average expression

We used expression data from Fly Atlas 2 (Krause et al. 2020). For each gene, expression values (measured as FPKM) were obtained from eight larval tissues and 24 adult tissues (12 per sex). The larval tissues were central nervous system, midgut, hindgut, tubule, fat body, salivary gland, trachea, and garland. The following 10 tissue types were used from both adult males and females: eye, brain, ganglion, crop, midgut, hindgut, tubule, heart, fat body, and rectal pad. In addition, testis and accessory glands were used from males and ovary and spermatheca from females. A “larval average” was calculated across the eight larval tissues and an “adult average” across the 24 adult tissues; these mean of these two values was used as a joint “larval-adult” average. The natural log of the larval-adult average was used for analyses.

### Tissue-specificity in expression

Using the expression data from Fly Atlas 2 (Krause et al. 2020), tissue specificity was calculated as: 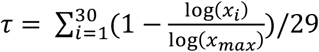 where *x*_*i*_ = 1 + the FPKM in tissue *i* (Yanai et al. 2005). The 30 tissues used were all of those listed above but excluding rectal pad. (In exploring the Fly Atlas 2, we noticed patterns suggesting that the male and female values may be reversed. Tissue specificity values are similar regardless of whether rectal pad is included or not.)

### Across-tissue expression profiles

Separately for adults of each sex, we calculated for each gene a vector of proportional expression across 11 tissues (as listed above, excluding rectal pad). The *i*^*th*^ element of the vector was expression in the *i*^*th*^ tissue (FPKM) divided by the sum of that gene’s expression values across all 11 tissues. Separately for each sex, we performed robust principal component analyses on the proportional data using *pcaCoDa* (Templ et al. 2011). The first three principal components (PCs) for each sex were used in downstream analyses. These PCs accounted for 79% and 71% of the variation in expression profiles in males and females, respectively. Tissue loadings for each PC are given in Table S2.

### Intersexual genetic correlation *r*_*mf*_

Following Fraïsse et al. (2019), we calculated the intersexual genetic correlation in gene expression from the data of Huang et al. (2015) who measured whole body gene expression of 185 genotypes from the Drosophila Reference Genome Panel. Using *MCMCglmm* (Hadfield 2010), for each gene we fit the model *z*_*ijk*_ = *µ* + *ϕ*_*jk*_ + *e*_*ijk*_ where *z*_*ijk*_ is the normalized expression of the *i*th replicate of sex *j* of genotype *k. µ* is the mean expression. *ϕ*_*jk*_ is the random genetic effect of sex *j* of genotype *k* and follows a bivariate Gaussian distribution characterized by sex-specific variances and the intersexual covariance. *e*_*ijk*_ is the residual. Models were fit using uninformative priors and run for a million iterations after a burn-in period of 10,000 iterations with a thinning interval of 1000. From the resulting MCMC posterior of sex-specific variances and intersexual covariances, a posterior of *r*_*mf*_ was created and the mean value used as the estimate of *r*_*mf*_. We only considered *r*_*mf*_ values for genes for which both sexes had genetic variances > 0.001 and statistically significant (*p* < 0.05) in at least one sex based on results of Huang et al. (2015; their Table S5). In the Supplementary Material we show another version of the main figure using only genes for which the genetic variance was statistically significant in both sexes; the patterns are very similar.

### Population Genomic Data

We obtained whole genome sequences for 205 haploid genotypes of the DPGP3 *Drosophila* population which were originally collected from a wild population in Siavonga, Zambia (Lack et al. 2015, 2016). FASTA files were obtained from the *Drosophila* Genome Nexus (DGN; Lack et al. 2015, 2016) website (https://www.johnpool.net/genomes.html) and were filtered to remove sites with evidence of high relatedness and recent admixture using Perl scripts provided on the DGN website. Following Sohail et al. (2017), we removed five genotypes that had extremely high or low numbers of variants. Additionally, we removed a single genotype (ZI28) for which the length of the X chromosome in the FASTA file differed from the other 204 genotypes. We then created a VCF file using the remaining 199 sequences with *SNP-sites* (v2.5.1; Page et al. 2016). Coordinates for the VCF file were converted to release 6 of the *Drosophila* reference genome using *UCSC liftOver* (Kent et al. 2002).

### Estimating nucleotide diversity and Tajima’s *D*

We obtained a GFF file for the *D. melanogaster* genome from Ensembl (https://useast.ensembl.org/Drosophila_melanogaster/) and scored each coding site in the genome as 4-fold (i.e., synonymous sites) or 0-fold (i.e., non-synonymous sites) degenerate across the *Drosophila* genome using a Python script (script available at https://github.com/tvkent/Degeneracy). We then parsed our VCF containing variant and invariant sites in the DPGP3 into two files that contained either only 4-fold sites or only 0-fold degenerate sites. After excluding sites with data from fewer than 100 lines and those with more than two alleles, we measured per site nucleotide diversity *π* = 2*p*(1 − *p*), where *p* is the estimated frequency of the reference base. Nonsynonymous diversity *π*_*N;*_ within a gene was measured as the mean nucleotide diversity at 0-fold degenerate sites within that gene and synonymous diversity *π*_*S*_ within a gene as the mean nucleotide diversity at 4-fold degenerate sites within each gene. Our analysis focuses on a measure of *relative* nonsynonymous diversity, 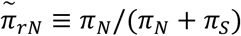. We use this rather than the more familiar ratio of *π*_*N*_/*π*_*S*_ because the latter has undesirable statistical properties, being highly sensitive to sampling variation in *π*_*S*_ and is undefined if *π*_*S*_ = 0. Our measure 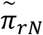_;_is bounded between 0 and 1 and will be similar to *π*_*N*_/*π*_*S*_ when *π*_*S*_ >> *π*_*N*_, as is expected for typical genes.

Tajima’s *D* (Tajima 1989) was estimated using 4-fold degenerate sites, accounting for the sample size and number of 4-fold sites (both invariant and variant) considered for each gene (Walsh and Lynch 2020, p. 303); sites with data from a fewer than 100 lines were not considered. When sites varied in sample size, we individually down sampled variant sites to the level of the site with the lowest sample size, then calculated Tajima’s *D*. Downsampling was performed 100 times per gene and the median value of Tajima’s *D* was used.

### Direction of Selection (*DoS*)

*DoS* (Stoletzki and Eyre-Walker 2011) is a metric that compares relative nonsynonymous divergence to relative nonsynonymous polymorphism; in both cases, it is relative to the sum of synonymous and nonsynonymous changes. Specifically, *DoS* = *D*_*N*_/(*D*_*N*_ + *D*_*S*_) – *P*_*N*_/(*P*_*N*_ + *P*_*S*_) where *D*_*N*_ and *P*_*N*_ are the numbers of nonsynonymous interspecific differences and intraspecific polymorphisms, respectively; *D*_*S*_ and *P*_*S*_ are defined analogously for synonymous changes. We used the *DoS* values reported by Fraïsse et al. (2019). They calculated divergence with respect to *D. simulans*. They calculated polymorphism from 197 haploid genomes of *D. melanogaster* genomes of the DPGP3. They excluded rare variants (minor allele frequency < 5%) to reduce the influence of deleterious variants on *DoS*.

### Statistical Analysis

For each of four evolutionary genetic properties of interest—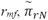, Tajima’s *D*, and *DoS*—we used linear models to examine whether among-gene variation in the metric was predicted by the following gene expression characteristics: SSS status, SBGE measured as log_2_FC, average expression, tissue specificity, as well as three male PCs and three female PCs capturing major axes of variation in among-tissue expression profiles. A quadratic term, SBGE^2^, was included to capture effects of sex bias irrespective of the direction of the bias. Genes with extreme sex-bias (|log_2_FC| > 5) were excluded from these analyses because visual inspection revealed that these genes had distinctly different values for several of the evolutionary genetic properties of interest (see Results). For the results presented in the main text, SSS status was coded as a numerical value: 2 (strong evidence of SSS: *p*_*adj*_ < 0.01), 1 (modest evidence: 0.2 < *p*_*adj*_ ≤ 0.01), or 0 (poor evidence: *p*_*adj*_ ≥ 0.2).

All models included an indicator variable of whether the gene was autosomal or X-linked. In addition, the models for each of the three population genomic metrics—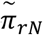, Tajima’s *D*, and *DoS*— included two other terms: log(recombination rate) and log(transcript length). Recombination is expected to affect population genetic patterns by mediating the effects of linked selection. Transcript length is also expected to affect linked selection as a given site in a long gene is expected to have more closely linked targets of selection than a site in a short gene. Recombination values came from Fraïsse et al. (2019), who obtained recombination measures from Chan et al. (2012).

The statistical support for model terms was assessed in two different ways. We obtained permutation-based *p*-values using the *lmp* function from the *lmperm* package (Wheeler and Torchiano, 2016). We used perm = “Prob”, Ca = 0.001, maxIter = 10^6^ and seq = FALSE for unique rather than sequential sum of squares. Second, we used the *Boot* function from the *boot* package (Canty and Ripley 2020) on each linear model, followed by *confint* to obtain 95% bootstrap confidence intervals of model terms. We specified method = “case” to sample over cases, which is considered a conservative approach. In almost all cases, the two methods gave consistent results, i.e., when terms were significant based on permutation *p*-values < 0.05, the bootstrap confidence intervals for the estimate did not overlap zero.

### Data from different sources

The data sets used here involves flies of different geographic origins. The expression data (Osada et al. 2017) comes from an F1 cross between a genotype from West Africa and genotypes from the DGRP (North America). We assume that these expression measures are reasonably representative of what is typical for the species and, most importantly, reasonably reflect the variation among genes in expression. That is, the within-gene variation (say, among genotypes or rearing conditions) is small compared to the among-gene variation that is our focus here. In support of this assumption, Zhang et al. (2007) found high correlations in SBGE across different species of *Drosophila* representing tens of millions of years of evolutionary divergence. The *r*_*mf*_ data come from the DGRP (Huang et al. 2015) and thereby represent a measure of genetic variation within a North American population. It seems plausible, though unverified, that *r*_*mf*_ values are similar in other populations. The population genetic metrics—Tajima’s *D*, 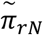, and *DoS*—are based on data from African lines. Because Africa is the ancestral range of *D. melanogaster*, it is preferable to use African lines (as others have, e.g., Fraisse et al. 2018) because the site frequency spectrum of North American samples will be affected by its colonization history.

## Acknowledgements

We wish to thank Tyler Kent for providing a script to identify *N*-fold degeneracy and Julia Kreiner, and James Santangelo for bioinformatics advice. Two anonymous reviewers provided helpful comments. Funding was provided in the form of an Ontario Graduate Scholarship to AS and an NSERC discovery grant to AFA.

