## Supplemental Material for "Two forms of sexual dimorphism in gene expression in *Drosophila melanogaster*: their coincidence and evolutionary genetics"

for

their coincidence and evolutionary genetics

by

Amardeep Singh and Aneil F. Agrawal

Table of Contents

**Supplemental Figures**

1. Histogram of $log_2FC$ values for body and head samples.
2. Proportion of genes exhibiting sex-specific splicing (SSS) considering only genes evaluated in both body and head.
3. The intersexual genetic correlation for expression ($r_{mf}$) with respect to SSS and SBGE. considering only genes with significant genetic variation in both sexes.
4. The divergence and polymorphism components of *DoS* with respect to SSS and SBGE.

**Supplemental Tables**

1. Results from GLM examining relationship between the probability of a gene being classified as SSS or not as function of SBGE.
2. Results for separate analyses of divergence and polymorphism components of *DoS*.
3. Tissue loadings for principal components of among-tissue expression profiles in males.
4. Tissue loadings for principal components of among-tissue expression profiles in females.

**SUPPLEMENTAL FIGURES**

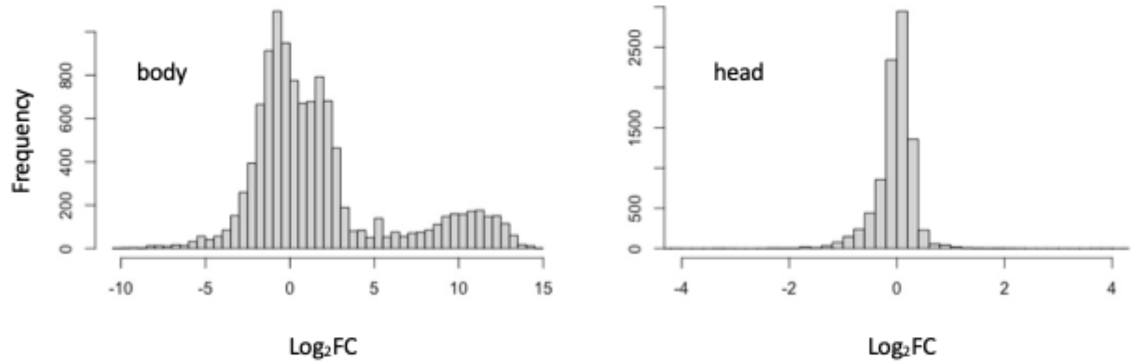


**Figure S1.** Histogram of log2FC values ("SBGE") for body (left) and head (right) samples. Negative/positive values indicate female/male bias, respectively.

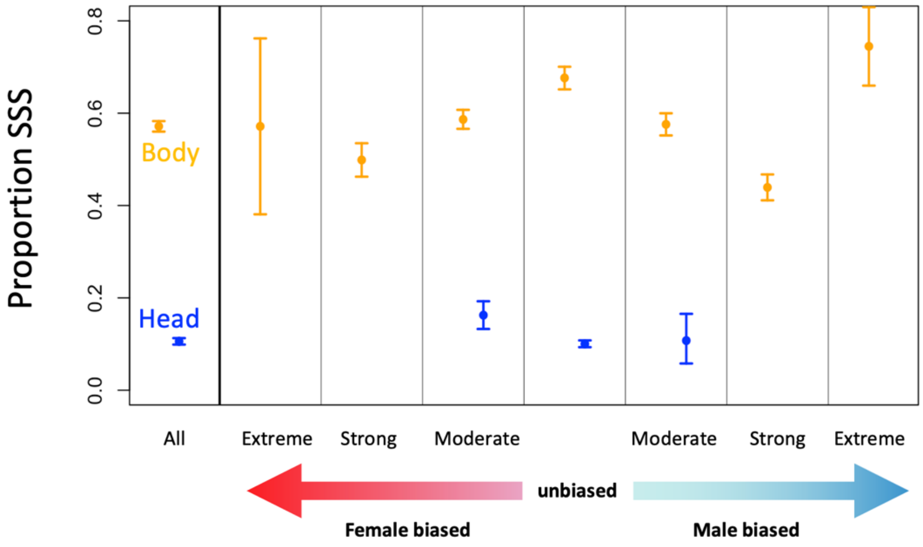


**Figure S2**. Proportion of genes exhibiting sex-specific splicing (SSS). In contrast to Figure 1 of the main text, only genes expressed and tested for SSS in both body and head are included here. The leftmost section shows results irrespective of SBGE; the remaining sections show results stratified by SBGE: extreme female bias ($log_2FC < -5$), strong female bias ($-5 \leq log_2FC < -2$ ), moderate female bias ($-2 \leq log_2FC < -0.5$), unbiased expression ($0.5 \leq log_2FC < 0.5$), moderate male bias ($0.5 \leq log_2FC < 2$), strong male bias ($2 \leq log_2FC < 5$), and extreme male bias ($log_2FC \geq 5$). Points in orange and blue represent reflect characterizations based on expression in body and head samples, respectively. Only three levels of SBGE are shown with respect to head because very few genes fall into the other levels. Error bars represent bootstrap 95% confidence intervals.

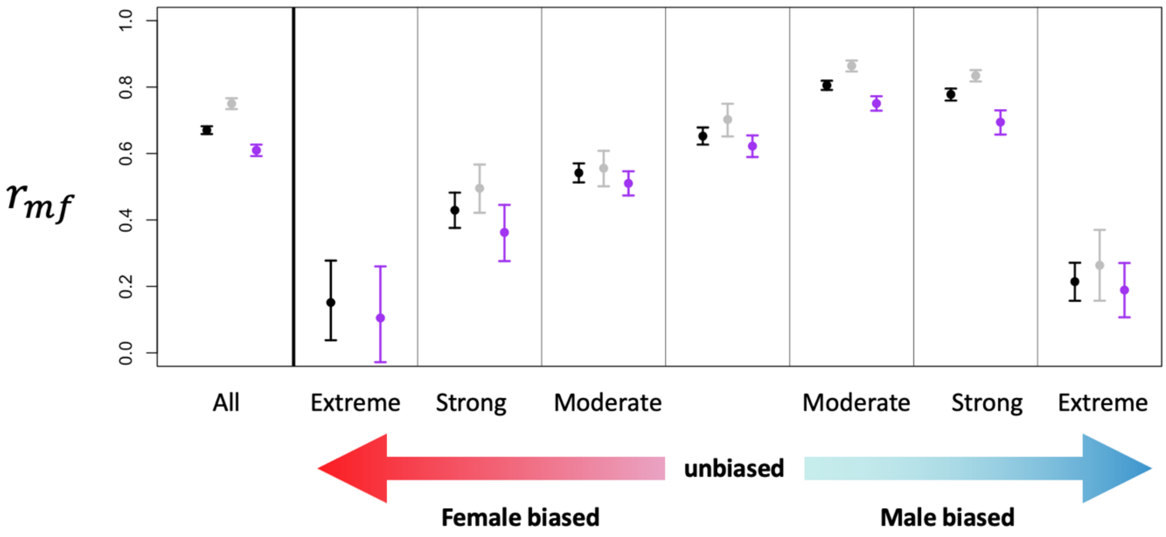


**Figure S3.** $r_{mf}$ with respect to SSS and SBGE including only genes for which Huang et al. (2015) found significant genetic variation in both sexes ($p < 0.05$). The leftmost section shows results irrespective of SBGE; the remaining sections show results stratified by SBGE: extreme female bias ($\log_2 FC < -5$), strong female bias ($-5 \leq \log_2 FC < -2$ ), moderate female bias ($-2 \leq \log_2 FC < -0.5$), unbiased expression ($0.5 \leq \log_2 FC < 0.5$), moderate male bias ($0.5 \leq \log_2 FC < 2$), strong male bias ($2 \leq \log_2 FC < 5$), and extreme male bias ($\log_2 FC \geq 5$). Points in grey and purple represent non-SSS and SSS genes, respectively; points in black are all genes irrespective of SSS status (including genes that were not tested for SSS so the black points represent more genes than the combined sum of genes represented by grey and purple points). Error bars are bootstrap 95% confidence intervals. The purple point for "extreme FB" is not shown because there are too few genes non-SSS genes (only 4).

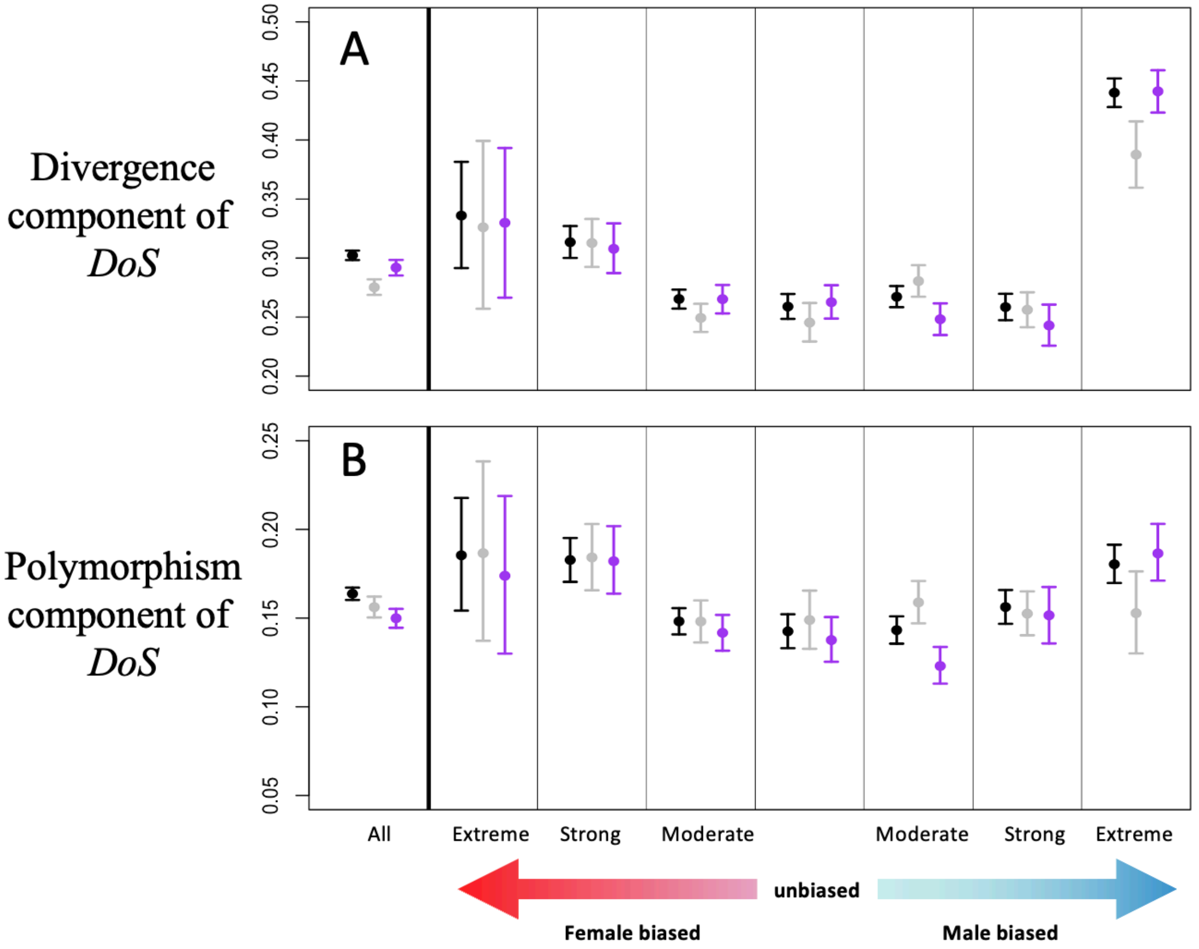


**Figure S4.** Divergence (**A**) and polymorphism (**B**) components of *DoS* with respect to SSS and SBGE. The leftmost section in each panel shows results irrespective of SBGE; the remaining sections show results stratified by SBGE. Points in grey and purple represent non-SSS and SSS genes, respectively; points in black are all genes irrespective of SSS status (including genes that were not tested for SSS so the black points represent more genes than the combined sum of genes represented by grey and purple points). Error bars are bootstrap 95% confidence intervals.

### SUPPLEMENTAL TABLES

**Table S1. Results from GLM examining relationship between the probability of a gene being classified as SSS or not as function of SBGE.**

| | Intercept | SBGE | $SBGE^2$ |
|---|---|---|---|
| Body | **0.30*****<br>[0.24, 0.36] | **-0.06*****<br>[-0.09, -0.04] | **-0.03*****<br>[-0.04, -0.02] |
| Head | **-2.23*****<br>[-2.31, -2.16] | **-0.30*****<br>[-0.58, -0.05] | **0.21*****<br>[0.02, 0.42] |

The numbers of genes used for the two analyses are $n$ = 7931 and 7754 for body and head, respectively. For these analyses, genes with extreme sex-bias ($log_2FC > |5|$) are excluded. The quadratic term, $SBGE^2$, is included to capture effects of sex bias that are independent of the direction of the bias. Estimates in bold have significant $p$-values ($z$-test): * $p < 0.05$, ** $p < 0.01$, ***$p < 0.001$. 95% bootstrap confidence intervals are shown in square brackets. Note estimates are on logit scale.

**Table S2. Results for separate analyses of divergence and polymorphism components of *DoS*.**

| Model Term | Divergence component of *DoS* | Polymorphism component of *DoS* |
|---|---|---|
| SSS | **0.007*** [0.002, 0.013] | -0.004 [-0.008, 0.001] |
| SBGE | **-0.012***** [-0.016, -0.008] | **-0.006**** [-0.009, -0.002] |
| $SBGE^2$ | 0.000 [-0.001, 0.002] | 0.000 [-0.001, 0.002] |
| Avg Expression | -0.005 [-0.010, 0.000] | **-0.016***** [-0.020, -0.002] |
| Tissue Specificity | **0.083***** [0.052, 0.114] | **0.076***** [0.050, 0.102] |
| M-PC1 | **-0.014***** [-0.018, -0.010] | -0.002 [-0.005, 0.001] |
| M-PC2 | 0.000 [-0.004, 0.005] | 0.001 [-0.002, 0.005] |
| M-PC3 | 0.000 [-0.003, 0.004] | **0.003*** [0.0002, 0.006] |
| F-PC1 | 0.000 [-0.004, 0.005] | 0.002 [-0.001, 0.006] |
| F-PC2 | **0.005**** [0.002, 0.009] | **0.004**** [0.001, 0.007] |
| F-PC3 | 0.001 [-0.004, 0.002] | -0.002 [-0.004, 0.001] |
| X-linkage | **-0.020***** [-0.027, -0.012] | **0.010***** [0.004, 0.015] |
| log(recombination) | 0.001 [-0.002, 0.004] | **-0.021***** [-0.024, -0.018] |
| log(gene length) | **-0.032***** [-0.040, -0.024] | -0.005 [-0.013, 0.002] |

For these analyses, genes with extreme sex-bias ($log_2FC > |5|$) are excluded. The quadratic term, $SBGE^2$, is included to capture effects of sex bias that are independent of the direction of the bias. Estimates in bold have significant permutation-based *p*-values: * $p < 0.05$, ** $p < 0.01$, ***$p < 0.001$. 95% bootstrap confidence intervals are shown in square brackets.

**Table S3. Tissue loadings for principal components of among-tissue expression profiles in males.**

| | M-PC1 | M-PC2 | M-PC3 | M-PC4 | M-PC5 | M-PC6 | M-PC7 | M-PC8 | M-PC9 | M-PC10 |
|---|---|---|---|---|---|---|---|---|---|---|
| Eye | 0.21 | -0.22 | 0.15 | 0.03 | 0.12 | -0.07 | 0.6 | 0.58 | 0.25 | -0.09 |
| Brain | 0.2 | -0.61 | 0.08 | 0.02 | -0.25 | -0.08 | -0.29 | -0.2 | -0.04 | -0.54 |
| Ganglion | 0.12 | -0.39 | 0.18 | 0.03 | -0.11 | 0.18 | -0.04 | -0.12 | -0.16 | 0.79 |
| Crop | 0.08 | 0.09 | 0.01 | 0.02 | 0.5 | 0.33 | 0.26 | -0.62 | 0.26 | -0.11 |
| Midgut | 0.23 | 0.18 | -0.31 | -0.46 | -0.07 | -0.66 | 0.12 | -0.21 | -0.04 | 0.14 |
| Hindgut | 0.25 | 0.12 | -0.21 | -0.18 | 0.49 | 0.2 | -0.55 | 0.4 | -0.13 | -0.01 |
| Tubule | -0.07 | 0.28 | -0.11 | -0.41 | -0.57 | 0.54 | 0.09 | 0.07 | 0.05 | -0.1 |
| Heart | -0.1 | 0.31 | 0.43 | 0.15 | 0.05 | -0.08 | 0.16 | -0.02 | -0.73 | -0.17 |
| Fat Body | -0.11 | 0.34 | 0.51 | 0.16 | -0.12 | -0.24 | -0.37 | 0.02 | 0.53 | 0.07 |
| Testis | -0.87 | -0.25 | -0.2 | -0.1 | 0.16 | -0.11 | -0.01 | 0.08 | 0.01 | 0.01 |
| Accessory Gland | 0.07 | 0.17 | -0.54 | 0.73 | -0.21 | -0.02 | 0.02 | 0.02 | -0.01 | 0.03 |
| % of Variance | 51.9 | 18.4 | 8.9 | 6.5 | 4.2 | 3.3 | 2.4 | 1.8 | 1.7 | 1.0 |

The bottom row shows the percent of variation accounted for each PC.

**Table S4. Tissue loadings for principal components of among-tissue expression profiles in females.**

| | F-PC1 | F-PC2 | F-PC3 | F-PC4 | F-PC5 | F-PC6 | F-PC7 | F-PC8 | F-PC9 | F-PC10 |
|---|---|---|---|---|---|---|---|---|---|---|
| Eye | -0.25 | 0.02 | 0.04 | 0.21 | 0.19 | -0.01 | 0.38 | 0.35 | 0.7 | -0.11 |
| Brain | -0.58 | -0.25 | -0.13 | -0.25 | 0 | -0.04 | -0.09 | -0.13 | -0.23 | -0.6 |
| Ganglion | -0.47 | -0.16 | -0.09 | -0.02 | 0.11 | 0 | -0.02 | -0.07 | -0.14 | 0.78 |
| Crop | 0.03 | 0.1 | -0.07 | 0.67 | -0.26 | 0.09 | 0.2 | 0.26 | -0.5 | -0.09 |
| Midgut | 0.23 | 0.29 | -0.38 | -0.2 | 0.04 | -0.76 | 0.04 | 0.08 | -0.06 | 0.02 |
| Hindgut | 0.07 | 0.21 | -0.24 | 0.31 | -0.21 | 0.14 | -0.31 | -0.65 | 0.36 | -0.02 |
| Tubule | 0.21 | 0.33 | -0.33 | -0.37 | 0.26 | 0.62 | 0.02 | 0.21 | -0.11 | -0.01 |
| Heart | 0.17 | -0.09 | 0.38 | 0.2 | 0.53 | -0.1 | -0.61 | 0.15 | -0.05 | -0.07 |
| Fat Body | 0.15 | 0.14 | 0.54 | -0.14 | 0.17 | -0.02 | 0.52 | -0.47 | -0.17 | -0.02 |
| Ovary | 0.47 | -0.78 | -0.14 | -0.09 | -0.16 | 0.05 | 0.12 | 0 | 0.09 | 0.03 |
| Virgin Spermatheca | -0.03 | 0.18 | 0.43 | -0.32 | -0.66 | 0.02 | -0.24 | 0.28 | 0.11 | 0.08 |
| % of Variance | 29.7 | 25.1 | 15.9 | 7.5 | 6.3 | 5.1 | 3.2 | 3.1 | 2.9 | 1.2 |

The bottom row shows the percent of variation accounted for each PC.
